# Thyroid hormone modulation during zebrafish development recapitulates evolved diversity in danionin jaw protrusion mechanics

**DOI:** 10.1101/418483

**Authors:** Demi Galindo, Elly Sweet, Zoey DeLeon, Mitchel Wagner, Adrian DeLeon, Casey Carter, Sarah McMenamin, W. James Cooper

## Abstract

One of three vertebrates belongs to a fish lineage for which protrusile jaws are a synapomorphy. Identifying the developmental determinants of protrusion ability will improve our understanding of an important area of evolutionary diversification. The high water viscosities experienced by tiny fish larvae inhibit the viability of protrusile jaws. In the zebrafish protrusion does not arise until after metamorphosis. Fish metamorphosis typically includes significant changes in trophic morphology, accompanies a shift in feeding niche and coincides with increased thyroid hormone production. We tested whether thyroid hormone affects the development of zebrafish feeding mechanics. We found that it affected all developmental stages examined, but that these effects were most pronounced after metamorphosis. Thyroid hormone levels affected the development of jaw morphology, feeding mechanics, shape variation and cranial ossification. Adult zebrafish utilize protrusile jaws, but an absence of thyroid hormone eliminated postmetamorphic remodeling of the premaxilla and the premaxillary structure that permits protrusion never formed. The premaxillae of late juvenile and adult zebrafish are similar to those found in the adults of other *Danio* species. Premaxillae from early juvenile zebrafish and hypothyroid adult zebrafish resemble those from adults in the genera *Danionella*, *Devario* and *Microdevario* that show little to no jaw protrusion.

## INTRODUCTION

Many if not most fishes undergo a metamorphosis during which their bodies are extensively remodeled (McMenamin and Parichy 2013). It is during metamorphosis and post-metamorphic development that most fishes acquire the behavioral and morphological characters that allow them to occupy their adult niches. Pre-metamorphic (larval) and post-metamorphic (juvenile and adult) fishes of the same species frequently live in different habitats and occupy disparate feeding niches (McCormick and Makey 1997; McCormick et al. 2002; Leis and McCormick 2006; McMenamin and Parichy 2013). The ecological diversification of adults in many fish lineages is therefore closely linked to evolutionary changes in the processes that shape their post-larval development. Most molecular developmental studies have focused on early development, particularly embryonic stages (Albertson and Yelick 2004; Parsons et al. 2010; Cooper et al. 2013; McMenamin and Parichy 2013), but if we are to understand the developmental changes that have permitted the adaptive diversification of adult fish feeding we need a better understanding of the controls of morphogenesis in late development.

Thyroid hormone (TH) signaling plays a major role in directing late-developmental skeletal remodeling in vertebrates, and TH stimulates metamorphosis or metamorphosis-like processes in many species (Das et al. 2006; Paris et al. 2010; Laudet 2011; McMenamin and Parichy 2013; Wojcicka et al. 2013). Thyroid hormone is known to play an important role in skull morphogenesis and multiple cranial malformations are associated with aberrant TH signaling (Hanken and Hall 1988; Hanken and Summers 1988; Hirano et al. 1995; Desjardin et al. 2014). The reshaping of multiple skull bones must be coordinated during the larva-to-juvenile transition for a fish to have a properly integrated mature skull. This need for integrated development should be particularly strong for fishes with complex, highly kinetic adult skulls in which motion is transferred through multiple linages. Hormones reach all body organs essentially simultaneously via circulating blood, and therefore have the potential to act as agents of developmental coordination that stimulate multiple organs, including different bones, to transform at the same time.

Protrusile jaws are an important evolutionary innovation in fish feeding, but the advantages of jaw protrusion may be limited to post-metamorphic fishes. Highly moveable skull linkages that allow the jaws to protrude forward from the face have evolved independently in at least six lineages of bony fishes and protrusion has been lost, gained, reduced and enhanced many times in these clades (Ferry-Graham et al. 2008; Staab et al. 2012; Wainwright et al. 2015). Two of these, Cypriniformes (∼3,200 species, including the zebrafish) and Acanthomorpha (∼17,000 species), have been particularly successful and together comprise more than one-third of living vertebrates (Yang et al. 2010; Staab et al. 2012; Near et al. 2013; Wainwright et al. 2015). Maximum jaw protrusion distance has been closely linked with diet (Cooper, Carter, et al. 2017) and an ability to rapidly transition between morphs capable of different degrees of protrusion appears to support diversification into different feeding niches (Ferry-Graham, Gibb, and Hernandez 2008; Cooper and Westneat 2009; Cooper et al. 2010; Staab et al. 2012; Wainwright et al. 2015).

Although protrusile jaws can confer multiple functional abilities in adults, particularly the enhancement of suction production via rapid expansion of the mouth cavity (*i.e*., buccal cavity; Konow and Bellwood 2005; Ferry-Graham, Gibb, and Hernandez 2008; Holzman et al. 2012; Staab et al. 2012), the relatively high water viscosities experienced by small aquatic organisms severely limits the utility of protrusile jaws in fish larvae (Hernandez 2000; Hernández et al. 2002; Yaniv et al. 2014). Small fishes live in a low Reynolds number environment in which viscous forces are greater than inertial forces (Hernandez 1995, 2000). Under these conditions protrusile jaws would be more likely to push food items away than to draw them into the mouth via suction. Jaw protrusion does not arise until after metamorphosis in the zebrafish (Hernández, Barresi, and Devoto 2002; McMenamin et al. 2017), and the fluid mechanics of feeding in very small fishes suggests that this may be true of all protrusile species unless their larvae reach unusually large sizes.

In addition to the known roles of TH in both vertebrate metamorphosis and cranial morphogenesis, changes to TH signaling may have been an important component of the diversification of the functional morphology of cypriniform feeding (Shkil et al. 2012; Shkil et al. 2015; Shkil and Smirnov 2015; McMenamin, Carter, and Cooper 2017). In order to better understand the controls of fish metamorphosis and the developmental determinants of cypriniform jaw protrusion ability we measured the effects of modulating TH levels on the development of the functional morphology of feeding in the zebrafish. Since anatomical variation and covariation strongly affect evolutionary potential (Martinez-Abadias et al. 2009; Cooper et al. 2011; Simon et al. 2016), we also determined whether TH levels affect developmental variation in cranial morphology and the shape covariation between different skull regions. To do this we collected morphological and functional data from a developmental range of zebrafish in which TH production was elevated (TH+), normal, or eliminated (TH-) and morphological data from nine additional species of danionin minnows (Danionini; Danioninae; Cyprindae) that exhibit extensive diversity in adult jaw protrusion ability. We tested the following predictions: 1) normal TH levels are required for the development of functional abilities important to adult zebrafish feeding; 2) elimination of TH production will reduce developmental variation in zebrafish head shape; 3) normal TH levels are required for the development of the wild-type pattern of covariation between different regions of the zebrafish skull; and 4) the functional morphology of jaw protrusion in adult TH-zebrafish closely resembles that in related minnows with limited protrusion abilities.

## MATERIALS AND METHODS

### Study System

We utilized three zebrafish lines to study the effects of TH on the development of their feeding biomechanics: 1) the transgenic line *Tg(tg:nVenus-2a-nfnB)*^*wp.rt8*^ in which the thyroid follicles can be chemically ablated; 2) the mutant line *opallus*^*b1071*^, hereafter *opallus*, which has a missense mutation in thyroid stimulating hormone receptor (*tshr*) that causes constitutive hyperthyroidism (*i.e*., elevated TH levels; McMenamin et al. 2014); and 3) the AB wild-type line (euthyroid, *i.e*., normal TH). Both the transgenic and the mutant lines originated in the AB line (McMenamin et al. 2014). All fish were maintained under standard conditions at 28° C and a 14-hour light/10-hour dark cycle under approved WSU IACUC protocol 04285.

Hypothyroid specimens (TH-) were produced via nitroreductase-mediated cell ablation of thyroid follicles in *Tg(tg:nVenus-2a-nfnB)*^*wp.rt8*^ larvae at 4 days post-fertilization (dpf) following McMenamin et al. (2014). Ablation was performed immediately after formation of the thyroid follicles so that they were rendered permanently incapable of hormone production. In brief, TH-fish were produced by placing specimens in a solution of 1% di-methyl sulfoxide (DMSO) and 10 mM metronidazole for 4 hours. Control *Tg(tg:nVenus-2a-nfnB)*^*wp.rt8*^ fish treated with 1% DMSO only for 4 hours will be referred to as “DMSO” hereafter.

All specimens were fed live paramecia from 5 dpf until they were large enough to feed on live *Artemia* (at ∼14 dpf). In order to provide more nutritionally complete *Artemia*, newly hatched brine shrimp were collected after 24 hours and fed for an additional 24 hours with an infusion of *Spirulina* sp. algae (RGcomplete, Reed Mariculture, Inc., Campbell, CA, USA). In order to prevent exposure to exogenous TH present in most prepared fish foods, all fish were fed exclusively with enriched *Artemia* after they were large enough to be fully weaned off of paramecia. All specimens were maintained in 9 L aquaria from 5 dpf onward on a recirculating system with carbon filters. Specimens from each treatment were sampled for kinematic and shape analyses at 8, 15, 30, 60 and 100 dpf.

### Kinematic Analyses

Fish were filmed in lateral view while feeding on either paramecia (8 and 15 dpf) or *Artemia* (30, 65 and 100 dpf). We analyzed kinematic data from 5 feeding strikes for each individual and examined 5 specimens of each age class from each treatment group. Feeding strikes were recorded at 500 frames s^-1^ using an Edgertronic monochrome high-speed video camera (Sanstreak Corp., San Jose, CA, USA). Kinematic analyses of feeding strikes were performed using the ImageJ software program (Schneider et al., 2012). We measured the following variables in each video frame of all feeding strikes (see Fig. 1A for reference): gape distance (linear distance between landmarks 1 and 3), gape angle (the angle created between landmarks 1, 2 and 3 with 2 as the apex), jaw protrusion (linear distance between landmarks 1 and 6 minus the minimum distance recorded between 1 and 6 for that strike), hyoid depression (linear distance between landmarks 4 and 5 minus the minimum distance recorded between 4 and 5 for that strike), and cranial angle (the angle created by the intersection of a line running along the dorsal edge of the head with a line running along the dorsal edge of the body, with landmark 6 denoting the point of head rotation). All measurements were made by the same researcher to minimize introduction of error. Gape distance, jaw protrusion, and hyoid depression were standardized by fish standard length. Maximum values for every variable were recorded for each feeding strike of every specimen. These maxima were used to calculate a mean value for each specimen. For each age class we used ANOVA to test for differences between treatment groups for the maximum value of each variable. When a significant difference was detected we used a Tukey’s honestly significant difference (HSD) test to determine which treatment groups were significantly different from each other.

**Figure 1.**
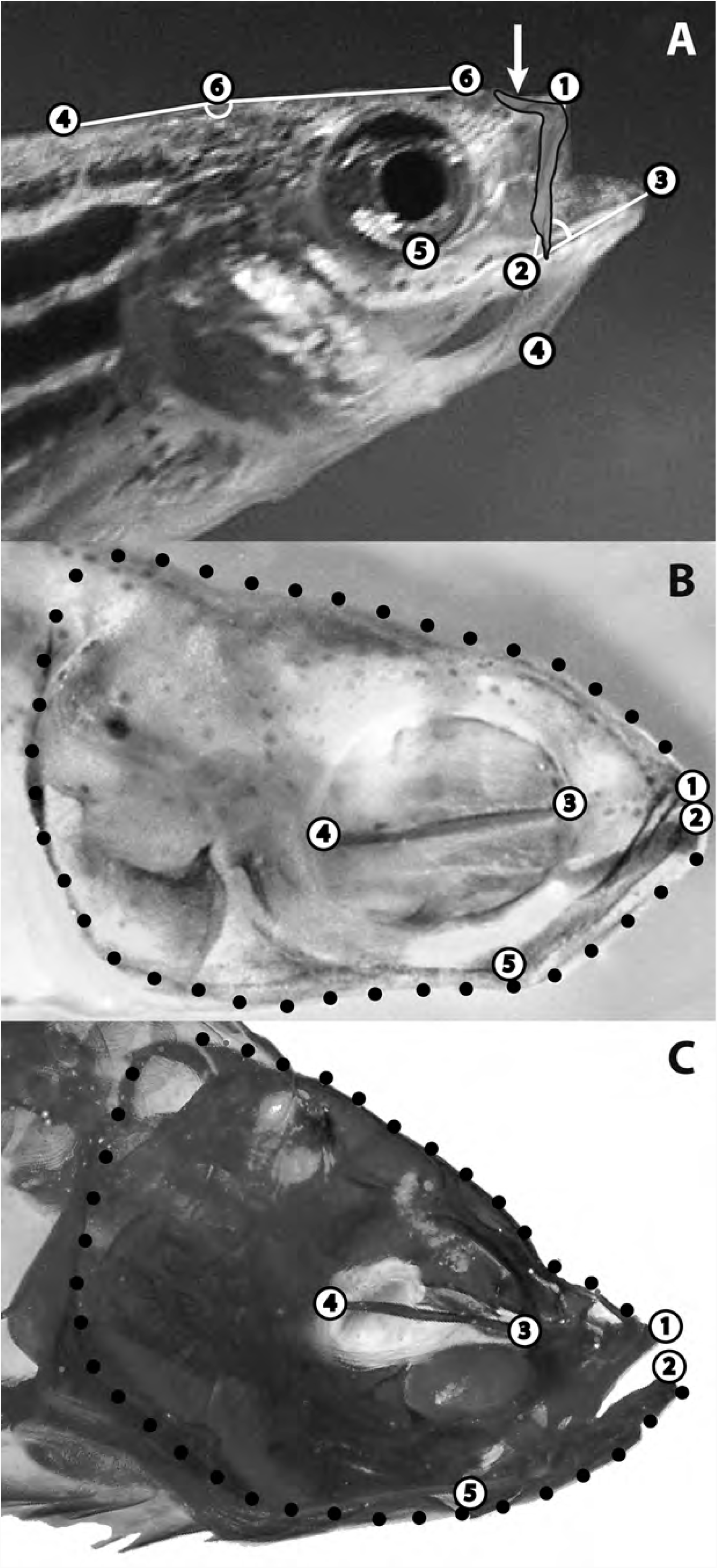
Landmarks and semi-landmarks used in analyses of movement and shape. A. Landmarks used in kinematic analyses with an image of the premaxillary bone of the upper jaw superimposed in correct anatomical position. Arrow indicates the ascending arm of the premaxilla. Landmarks: 1) Anterior tip of the upper jaw; 2) corner of the mouth; 3) anterior tip of the lower jaw; 4) anterior tip of the hyoid; 5) ventral-most point of the orbit; and 6) apex of the angle used to measure head rotation during cranial elevation (the dorsal surface of the head anterior to this point rotated upward during cranial elevation when feeding, while the dorsal surface of the trunk posterior to this point did not). B. Anatomical landmarks and semi-landmarks used in shape analyses of all specimens at all ages sampled shown on a larval zebrafish. Landmarks: 1) anterior tip of the premaxilla in the upper jaw; 2) anterior tip of the dentary bone in the lower jaw; 3) junction of the parasphenoid with the anterior wall of the orbit; 4) junction of the parasphenoid with the posterior wall of the orbit; and 5) lower jaw joint (articular-quadrate joint). Black circles indicate semi-landmarks evenly spaced between LM 1 and 2 in order to capture overall head shape. C. Anatomical landmarks and semi-landmarks used in shape analyses of all specimens at all ages sampled shown on an adult zebrafish.

### Analyses of Morphological Variation and Covariation

The specimens used in feeding trials were also used for morphological analyses. Euthanized specimens were fixed in formalin and then stepped over into 70% ethanol. They were then cleared and stained for bone and cartilage. The smaller specimens (8-30 dpf) were processed according to Walker and Kimmel’s (2006) acid-free staining protocol. Larger specimens (65 and 100 dpf) were cleared and stained according to Potthoff (1984). Specimens were then stepped into 80% glycerol and photographed in lateral view using an Olympus DP25 digital camera interfaced with an Olympus SZ61 dissecting microscope.

The program tpsDIG2 (http://life.bio.sunysb.edu/morph/) was used to place landmarks (LM) and semi-landmarks (semi-LM) on digital images of fish heads (Fig 1B,C). We chose skeletal LM that are present at all of the developmental stages examined. We did not place LM or Semi-LM were used to capture the shape of curved surfaces LM (Fig. 1B and C). The program tpsrelW (http://life.bio.sunysb.edu/morph/) was then used to superimpose semi-LM using a chord-distance (Procrustes distance) based ‘sliders’ method and to remove size and orientation differences from LM and semi-LM position data via Procrustes transformations. Pooled shape data from all specimens of all ages in each hormonal treatment group (N=25 for each treatment) were used to: 1) test for differences in head shape variation over the course of development; and 2) test for differences in the patterns of covariation between LM and semi-LM locations over the course of development.

Head shape variation was quantified by calculating the Foote disparity value for each treatment group (Foote 1993). We used a permutation procedure (2,000 iterations) to test for differences in disparity between pairs of datasets. If actual differences in shape disparity values were greater than the upper bound of a 95% confidence interval calculated via permutation, then the disparity values were considered to be significantly different. We used DisparityBox, which is an analytical tool available within the PCAGen8 program, to perform these calculations.

We performed principal components analysis (PCA) of LM and semi-LM positions using the program PCAGen8. These PCAs utilized covariation matrices that capture patterns of positional covariation. In order to determine if patterns of covariation were significantly different between the hormonal treatments we conducted pairwise comparisons of PC axis orientations using a bootstrapping procedure (4900 sets).

Although the different PC axes derived from the same data are orthogonal to each other, they are not independent, since only those aspects of covariation that were not associated with PC1 can be used to define subsequent axes. It therefore impossible to compare the orientations of PC axes subsequent to PC1 individually. All analyses that involved multiple axes determined whether the alignments of planes (2 axes) or multi-dimensional hyperplanes (≥3 axes) were significantly different.

### Premaxilla Shape Analyses

We also compared developmental variation in the premaxillary shape of AB and TH-zebrafish to the variation in adult premaxillary shape that has evolved among other members of the cyprinid tribe Danionini (*sensu* Tang et al. 2010). We obtained specimens of the following fishes through the pet trade: *Danio albolineatus*, *D. erythromicron*, *D. feegradei*, *D. kyathit*, *D. nigrofasciatus*, *Danionella translucida*, *Devario aequipinnatus*, *Devario maetaengensis*, and *Microdevario kubotai*. Adult specimens (2-4 per species) were cleared and stained following Potthoff (1984). AB zebrafish at 35, 65 and 100 dpf were also cleared and stained (5 specimens per age class). We used 35 dpf specimens instead of 30 dpf fish because some 30 dpf specimens did not have well-ossified premaxillae. Premaxillae were removed from all specimens after clearing and staining and then photographed as descried above. Anatomical LM and semi-LM (Fig. 4 B,C) were used to quantify premaxillary shape.

For all shape analyses we used the program CoordGen8 to transform tps formatted LM and semi-LM coordinate data into the format utilized by Mac OS (Apple Inc.) versions of the IMP-8 series of programs. The IMP-8 programs CoordGen8, PCAGen8, TwoGroup8 and PCAGen8 were developed by David Sheets and are available for download at: http://www3.canisius.edu/∼sheets/IMP8.htm.

## RESULTS

All specimens possessed largely cartilaginous crania at 8 and 15 dpf. Skulls were mostly, but not completely, ossified by 30 dpf in specimens from all treatment groups except TH- (Fig. 2). We will refer to 8 and 15 dpf specimens as “pre-metamorphic”, 30 dpf specimens as “mid-metamorphic” and 65 and 100 dpf specimens as “post-metamorphic.” Euthyroid and TH+ fish had fully ossified skulls by 65 dpf, while TH- specimens retained cartilaginous regions in the calvarium (skull vault) at this stage and in some TH- fish this region failed to ossify by 100 dpf (Fig. 2). Fish standard lengths did not significantly differ between the AB, DMSO, and TH+ treatments at any developmental stage (Fig. 3). Standard lengths were not significantly different for any treatment at 8, 15 and 30 dpf, but 65 and 100 dpf TH- fish were shorter than those of the same age from other treatments (Fig. 3). The lower jaws of TH+ specimens exhibited incipient enlargement by 30 dpf that became pronounced by 65 dpf (Fig. 2,3). Lower jaw morphology was normal in TH-specimens (Fig. 2,3). The maxillary and premaxillary bones of the upper jaw were smaller in 65 and 100 dpf TH- fish relative to euthyroid and TH+ specimens (Fig. 2). The ascending arm of the premaxilla, which determines maximum jaw protrusion distance (Fig. 1), was reduced in TH- fish (Fig. 2, 4). Upper jaw morphology was normal in TH+ specimens. Head shape differences between TH treatments were present at all developmental stages (Fig. S1). By 100 dpf TH+ and TH- head shapes were distinct from each other and from those of both AB and DMSO specimens. AB and DMSO specimens had the most similar head shapes at 100 dpf (Fig. S1).

**Figure 2.**
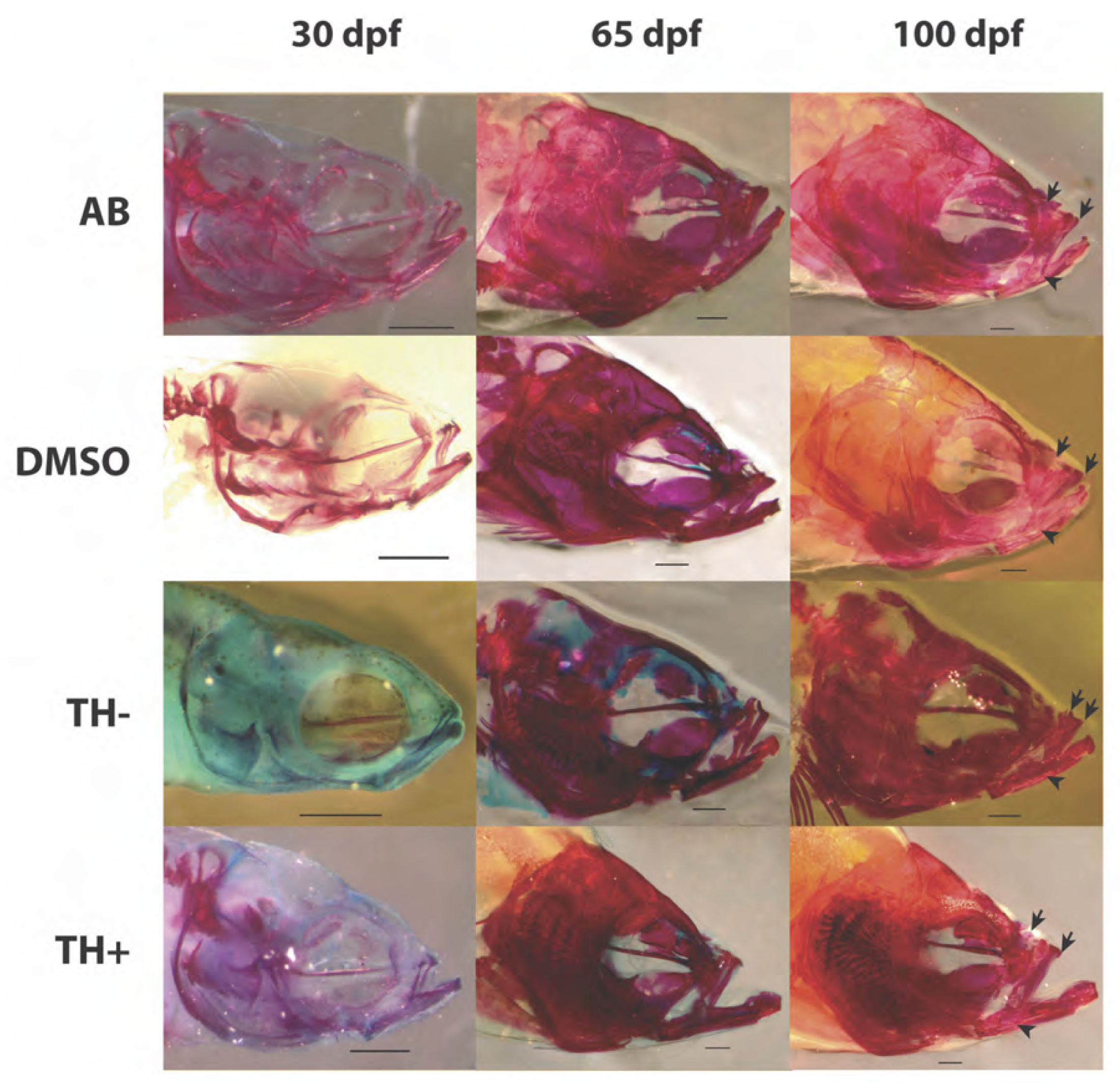
Representative cleared and stained specimens of post-metamorphic specimens from all treatment groups. Blue coloration indicates cartilage stained by alcian blue. Red indicates bone stained by alizarin red. Arrows represent the anterio-dorsal and posterio-dorsal edges of the ascending arm of the premaxillary bone. Arrowheads indicate the posterio-ventral tip of the dentigerous arm of the premaxilla. Scale bars = 1 mm. TH-specimens exhibited delayed cranial ossification. In many TH-specimens the calvarium (skull vault) never fully ossified. Both the overall size of the premaxilla and the length of its ascending arm relative to the length of its dentigerous arm were reduced in TH- fish. TH+ fish exhibited hypertrophied lower jaws.

**Figure 3.**
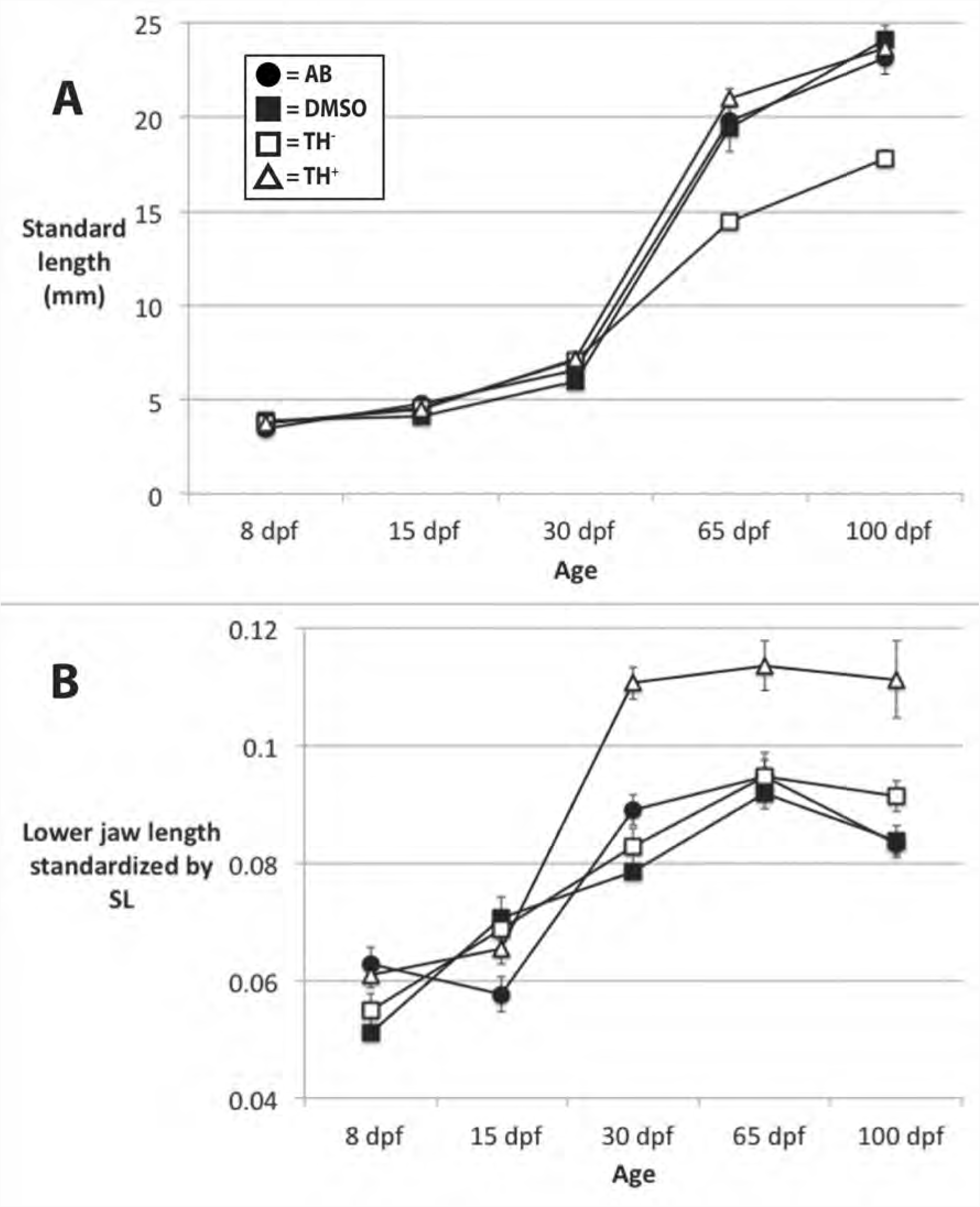
Comparisons of anatomical growth among treatment groups. Mean sizes with standard error bars are given for all treatments at each age sampled. Key to symbols in panel A. A. All treatment groups exhibited similar increases in body length until 30 dpf, after which TH- fish exhibited lower growth rates. B. All treatment groups exhibited similar increases in the relative length of the lower jaw until 15 dpf. After 15 dpf the lower jaws of TH+ fish grew much more quickly than those of the other treatment groups and then stabilized at a larger relative size.

**Figure 4.**
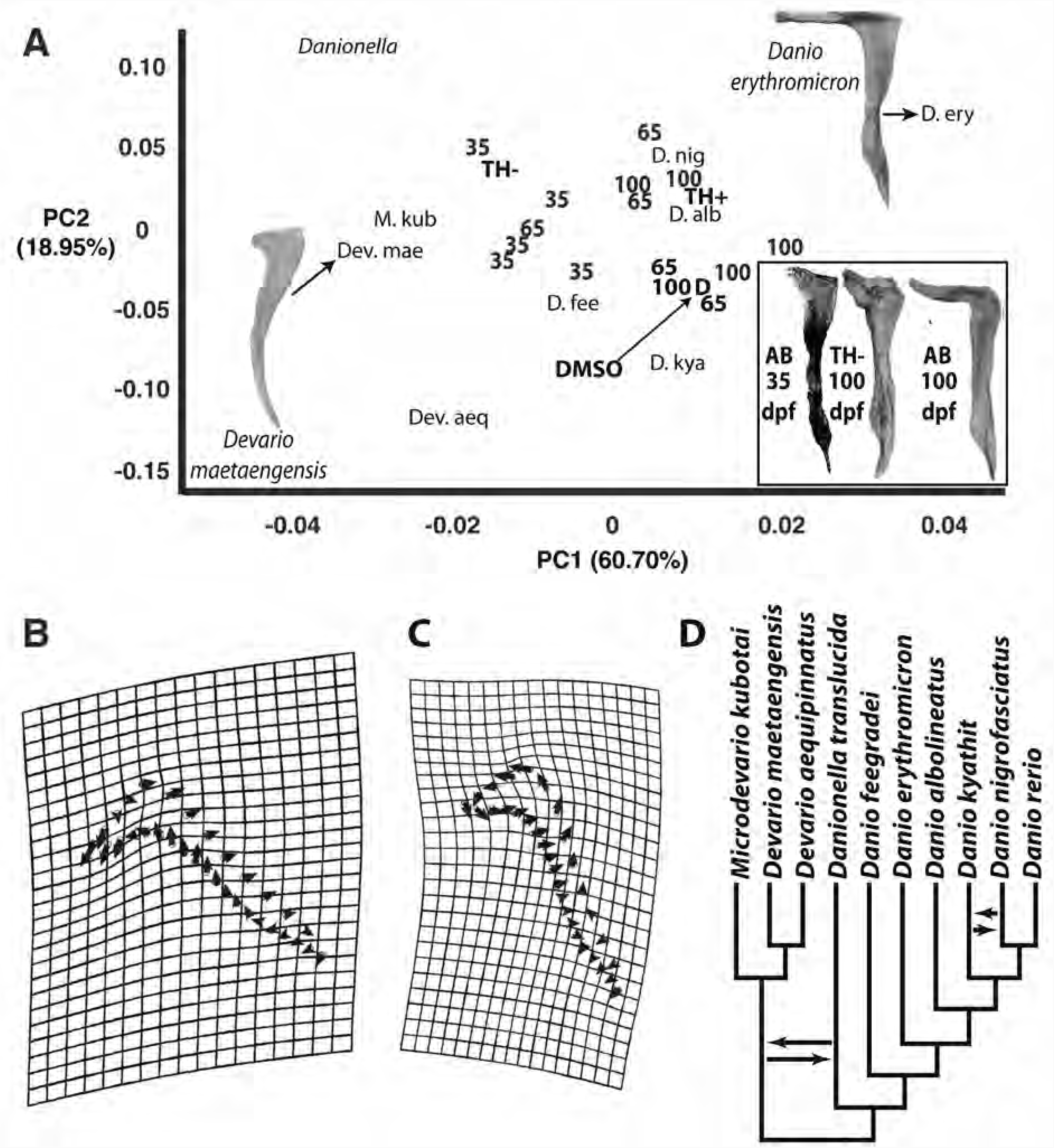
Comparisons of premaxillary morphology. A. Principal component score plot derived from coordinate-based analyses of premaxilla shape in the following specimens: 1) 35, 65 and 100 dpf AB zebrafish; 2) 100 dpf DMSO, TH- and TH+ zebrafish; and 3) adults of nine additional species of danionin minnows. The difference in premaxillary shape between *Devario maetaengensis* and *Danio erythromicron* exemplify the shape diversity explained by PC 1. PC1 is strongly associated with the length of the ascending arm relative to the length of the dentigerous arm. Numbers indicate the distributions of 5 specimens of AB zebrafish from each of the three ages sampled (numbers correspond to dpf). The location of the Procrustes mean premaxillary shapes of TH+ and TH- specimens are indicated by their respective symbols. The location of the Procrustes mean premaxillary shape of DMSO specimens is indicated by “D”. The location of Procrustes mean shapes of the nine non-zebrafish premaxillae are indicated by an abbreviation of the scientific name of each species (see panel D for full species names), except for *Danionella translucida*, where the complete genus name is used. Images of premaxilla shapes of particular interest are inserted. B. Deformation grid and vector plot that shows the shape variation associated with PC1. C. Deformation grid and vector plot that shows the shape variation associated with PC2. D. Phylogenetic relationships of the 10 species whose premaxillae were compared. The phylogeny depicted is taken from Tang et al. (2010). Arrows indicate branch positions that are swapped in the relationships reported by McCluskey and Postlethwait (2015), who did not examine *Devario maetaengensis*.

### Kinematic analyses

There were more significant kinematic differences between treatment groups after metamorphosis than before metamorphosis (Table 1). Four kinematic variables could be measured at all of the developmental stages examined (cranial elevation, gape angle, gape distance and hyoid depression). At each developmental stage these four variables were compared among all treatment groups (6 comparisons per variable). In regard to these 4 variables there were therefore 48 comparisons of kinematic performance both before (8 dpf and 15 dpf) and after (65 dpf and 100 dpf) metamorphosis. Of the 48 pre-metamorphic comparisons, 13 of them exhibited significant differences (27.1 %), while there were significant differences between 20 of the 48 post-metamorphic comparisons (41.7%; Table 1). Treatment groups were most similar in regard to cranial elevation (6 significant differences total) and most different in regard to hyoid depression (12 significant differences total; Table 1). Fish from the euthyroid treatments (AB and DMSO) showed the most similarity in cranial movement, with only five significant kinematic differences between these two treatments (all developmental stages combined; Table 1). Fish from the TH- treatment exhibited the most limited range of cranial element motion during feeding (Table 1; Fig. 4; Fig. S2). There were also 30 significant kinematic differences (all developmental stages combined) between TH- specimens and all other treatment groups, which was the highest number for any of the treatment groups (Table 1). Taken together, these results support our first prediction that normal TH levels are required for the development of functional abilities important to adult zebrafish feeding. These data support our first prediction that some minimum level of TH is necessary for the normal development of zebrafish feeding biomechanics.

**Table 1.** ANOVA and Tukey’s HSD results (F-statistic above p-value) for comparisons of kinematic variables among treatment groups for each developmental stage sampled.

|  | Gape Distance | Gape Angle | Hyoid Depression | Cranial Elevation | Jaw Protrusion |
| --- | --- | --- | --- | --- | --- |
| 8 dpf ANOVA | 7.027<br>**0.003** | 21.161<br>**<0.001** | 10.327<br>**<0.001** | 5.594<br>**0.008** | NA |
| 8 dpf Tukey's HSD | AB v TH<sup>-</sup><br>**0.004**<br>AB v TH<sup>+</sup><br>**0.010** | AB v DMSO<br>**<0.001**<br>AB v TH<sup>-</sup><br>**<0.001**<br>AB v TH<sup>+</sup><br>**<0.001** | AB v TH<sup>-</sup><br>**0.007**<br>AB v TH<sup>+</sup><br>**0.002**<br>DMSO v TH<sup>-</sup><br>**0.023**<br>DMSO v TH<sup>+</sup><br>**0.006** | DMSO v TH<sup>+</sup><br>**0.005** | NA |
| 15 dpf ANOVA | 2.041<br>0.149 | 10.090<br>**<0.001** | 4.776<br>**0.015** | 3.207<br>0.051 | NA |
| 15 dpf Tukey's HSD |  | AB v TH<sup>-</sup><br>**0.001**<br>AB v TH<sup>+</sup><br>**<0.001** | AB v TH<sup>-</sup><br>**0.013** |  | NA |
| 30 dpf ANOVA | 11.500<br>**<0.001** | 5.168<br>**0.011** | 12.719<br>**<0.001** | 3.688<br>**0.034** | NA |
| 30 dpf Tukey's HSD | AB v TH<sup>+</sup><br>**<0.001**<br>DMSO v TH<sup>+</sup><br>**0.002**<br>TH<sup>-</sup> v TH<sup>+</sup><br>**0.001** | DMSO v TH<sup>+</sup><br>**0.020**<br>TH<sup>-</sup> v TH<sup>+</sup><br>**0.027** | AB v DMSO<br>**0.003**<br>AB v TH<sup>-</sup><br>**0.011**<br>DMSO v TH<sup>+</sup><br>**<0.001**<br>TH<sup>-</sup> v TH<sup>+</sup><br>**0.002** | TH<sup>-</sup> v TH<sup>+</sup><br>**0.027** | NA |
| 65 dpf ANOVA | 20.110<br>**<0.001** | 10.530<br>**<0.001** | 0.370<br>0.776 | 19.730<br>**<0.001** | 15.839<br>**<0.001** |
| 65 dpf Tukey's HSD | AB v TH<sup>-</sup><br>**<0.001**<br>DMSO v TH<sup>-</sup><br>**<0.001**<br>TH<sup>-</sup> v TH<sup>+</sup><br>**<0.001** | AB v TH<sup>-</sup><br>**<0.001**<br>DMSO v TH<sup>-</sup><br>**0.012** |  | AB v DMSO<br>**<0.001**<br>AB v TH<sup>-</sup><br>**<0.001**<br>DMSO v TH<sup>+</sup><br>**<0.001**<br>TH<sup>-</sup> v TH<sup>+</sup><br>**0.001** | AB v TH<sup>-</sup><br>**<0.001**<br>DMSO v TH<sup>-</sup><br>**0.001**<br>TH<sup>-</sup> v TH<sup>+</sup><br>**<0.001** |
| 100 dpf ANOVA | 25.327<br>**<0.001** | 11.779<br>**<0.001** | 11.295<br>**<0.001** | 0.516<br>0.677 | 29.136<br>**<0.001** |
| 100 dpf Tukey's HSD | AB v TH<sup>-</sup><br>**<0.001**<br>AB v TH<sup>+</sup><br>**0.003**<br>DMSO v TH<sup>-</sup><br>**<0.001**<br>TH<sup>-</sup> v TH<sup>+</sup><br>**<0.001** | AB v DMSO<br>**0.047**<br>AB v TH<sup>-</sup><br>**<0.001**<br>AB v TH<sup>+</sup><br>**0.013**<br>DMSO v TH<sup>-</sup><br>**0.039** | AB v DMSO<br>**<0.015**<br>DMSO v TH<sup>+</sup><br>**<0.001**<br>TH<sup>-</sup> v TH<sup>+</sup><br>**0.003** |  | AB v TH<sup>-</sup><br>**<0.001**<br>DMSO v TH<sup>-</sup><br>**<0.001**<br>TH<sup>-</sup> v TH<sup>+</sup><br>**<0.001** |

Significant p-values from ANOVA in bold/shaded. Only significant post-hoc comparisons are listed.

Maximum gape distance occurred at or near time 0 (the point at which food is enveloped by the mouth) for all treatment groups throughout development (Figure 5A, B; Fig. S2). Hypothyroid zebrafish exhibited low gape distances at 8 and 100 dpf (Fig. 5A, B; Fig. S2). Maximum gape angle occurred at or near time 0 for all treatment groups throughout development (Fig. 5C, D; Fig. S2). AB fish exhibited high gape angles throughout development (Fig. 5C, D; Fig. S2), while TH- fish exhibited low gape angles throughout development (Fig. 5A, B; Fig. S2). Maximum hypoid depression occurred after time 0 for all treatment groups throughout development (Fig. 5E, F; Fig. S2). Hypothyroid fish exhibited low levels of hyoid depression throughout development until 100 dpf, when they displayed a high degree of hyoid depression (Fig. 5E, F; Fig. S2). Maximum cranial elevation occurred at or immediately after time 0 throughout development (Fig. 5G, H; Fig. S2). Hypothyroid fish exhibited low levels of cranial elevation throughout development (Fig. 5G, H; Fig. S2). Measureable upper jaw protrusion was not observed prior to 30 dpf in any treatment group, but a small number of euthyroid and TH+ specimens exhibited marginal upper jaw movement at this stage. Specimens from all treatment groups except TH- exhibited protrusile upper jaws by 65 dpf. Hypothyroid fish exhibited low levels of jaw protrusion at both 65 and 100 dpf and were significantly different from all other treatment groups at both stages (Table 1; Fig. 5I, J).

**Figure 5.**
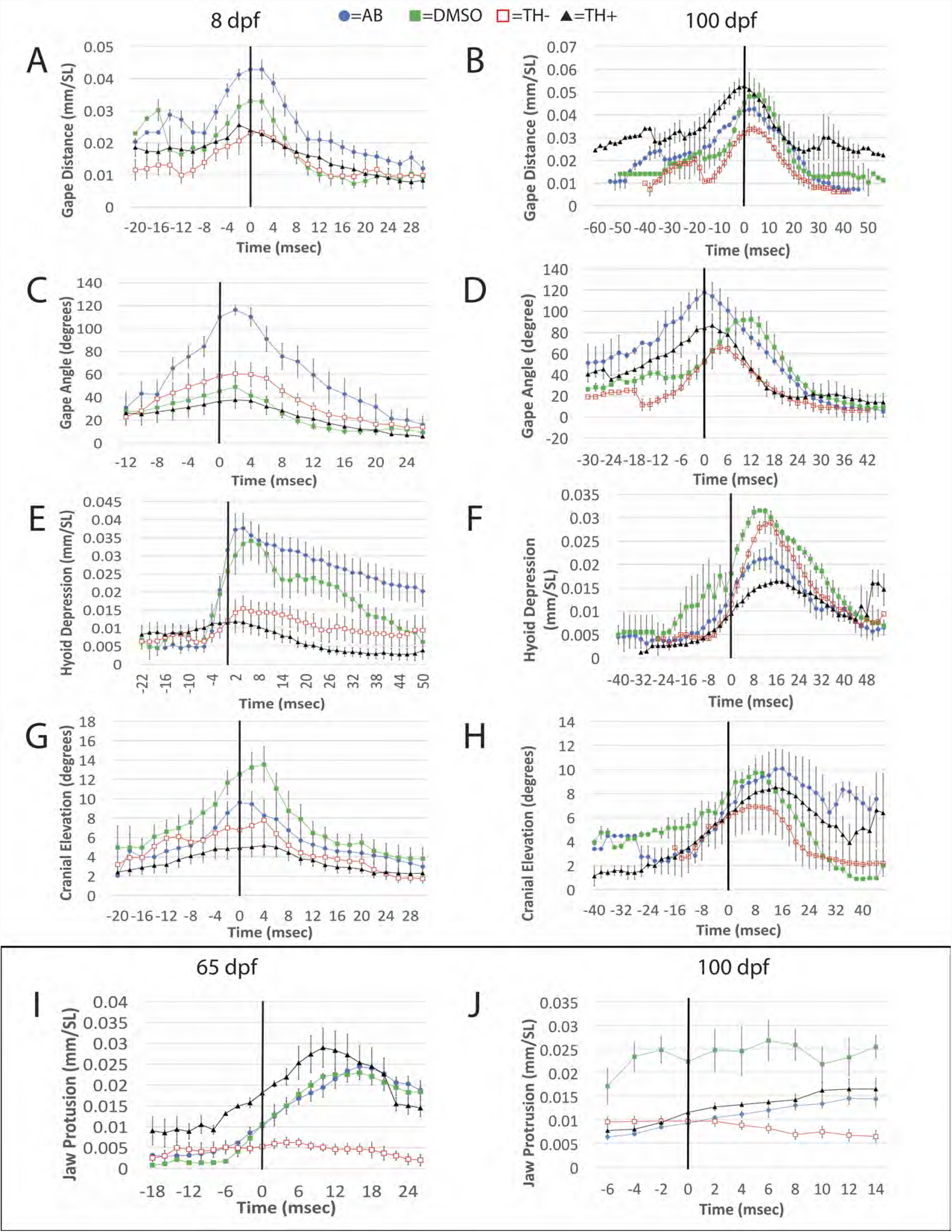
Comparisons of cranial movements during feeding among the four TH treatment groups. In all cases time zero represents the time point when live prey (paramecia at 8 dpf and brine shrimp nauplii at 65 and 100 dpf) passed the tips of the upper and lower jaws as they were being engulfed. Units and standardizations are given in parentheses in each case. SL = standard length. A-H. Plots for gape distance (A, B), gape angle (C, D), hyoid depression (E, F) and cranial elevation (G, H) for specimen of ages 8 and 100 dpf, respectively (kinematic plots for these four variables at 15, 30 and 65 dpf are presented in Fig. S2). I. Jaw protrusion at 65 dpf. J. Jaw protrusion at 100 dpf.

### Shape analyses

Hypothryoid fish showed significantly less developmental variation in head shape than any other treatment group (Table 2). Hyperthyroid fish showed significantly greater developmental variation in head shape than both TH- and DMSO specimens. These results strongly support our second prediction that elimination of TH production will reduce developmental variation in zebrafish head shape.

**Table 2.** Shape disparity comparisons for head morphology.

|  | AB | DMSO | TH<sup>-</sup> |
| --- | --- | --- | --- |
| AB<br>FD=0.00622046 | NA |  |  |
| DMSO<br>FD=0.00593240 | DD=0.000288<br>UBCI=0.00132652 | NA |  |
| TH<sup>-</sup><br>FD=0.00410838 | **DD=0.002112 (AB)**<br>UBCI=0.00139538 | **DD=0.001824 (DMSO)**<br>UBCI=0.00134556 | NA |
| TH<sup>+</sup><br>FD=0.00780772 | DD=0.001587<br>UBCI=0.00182215 | **DD=0.001875 (TH<sup>+</sup>)**<br>UBCI=0.00174844 | **DD=0.003699 (TH<sup>+</sup>)**<br>UBCI=0.00188894 |

FD=Foot disparity. DD=difference in the disparities of the treatments compared. UBCI=upper bound of 95% CI. Disparities were significant when DD > UBCI. Treatments with higher disparities are indicated (bold/shaded) when significant.

All treatments had head shape PC1 orientations that were not significantly different from parallel (Table 3). The PC2 axes of TH- and TH+ specimens were not parallel, but all other one-to-one comparisons between treatment groups revealed no differences in cranial shape space orientation for the first four PC axes (Table 3). We therefore reject our third prediction that normal TH levels are required for the development of the wild-type pattern of covariation between different regions of the zebrafish skull.

**Table 3.** Shape space orientation comparisons for head and premaxilla morphology.

| PC1 | Angle (degrees) | 95% CI |
| --- | --- | --- |
| AB vs. DMSO | 21.78 | 16.35-31.81 |
| AB vs. TH<sup>-</sup> | 34.73 | 29.24-44.89 |
| AB vs. TH<sup>+</sup> | 23.71 | 19.65-31.17 |
| DMSO vs. TH<sup>-</sup> | 18.26 | 15.50-30.77 |
| DMSO vs. TH<sup>+</sup> | 17.55 | 15.99-24.94 |
| TH<sup>-</sup> vs. TH<sup>+</sup> | 19.13 | 15.92-27.90 |
| AB pramaxilla development vs. Danionini premaxillae diversity | 33.34 | 25.76-64.45 |
| PC1-PC2 | Angle (degrees) | 95% CI |
| AB vs. DMSO | 61.51 | 50.06-85.52 |
| AB vs. TH<sup>-</sup> | 88.02 | 46.32-94.51 |
| AB vs. TH<sup>+</sup> | 86.73 | 47.20-91.12 |
| DMSO vs. TH<sup>-</sup> | 65.94 | 40.42-90.94 |
| DMSO vs. TH<sup>+</sup> | 74.15 | 42.80-91.19 |
| TH<sup>-</sup> vs. TH<sup>+</sup> | **29.86** | 34.57-89.13 |
| AB pramaxilla development vs. Danionini premaxillae diversity | 52.15 | 38.73-84.78 |
| PC1-PC3 | Angle (degrees) | 95% CI |
| AB vs. DMSO | 80.03 | 73.32-96.61 |
| AB vs. TH<sup>-</sup> | 65.80 | 55.39-102.35 |
| AB vs. TH<sup>+</sup> | 59.58 | 53.65-86.61 |
| DMSO vs. TH<sup>-</sup> | 84.21 | 67.73-102.18 |
| DMSO vs. TH<sup>+</sup> | 77.30 | 65.46-100.39 |
| AB pramaxilla development vs. Danionini premaxillae diversity | 52.83 | 50.17-96.84 |
| PC1-PC4 | Angle (degrees) | 95% CI |
| AB vs. DMSO | 101.83 | 84.63-116.01 |
| AB vs. TH<sup>-</sup> | 93.18 | 71.75-109.74 |
| AB vs. TH<sup>+</sup> | 77.48 | 66.79-106.49 |
| DMSO vs. TH<sup>-</sup> | 86.38 | 74.52-111.07 |
| DMSO vs. TH<sup>+</sup> | 97.36 | 77.98-117.97 |
| AB pramaxilla development vs. Danionini premaxillae diversity | 79.00 | 74.07-109.85 |

The observed angle between the shape spaces is given for each comparison. 95% confidence intervals for this angle were calculated by bootstrapping the data from both groups (700 bootstraps). Significant differences in bold/shaded. The TH<sup>-</sup> and TH<sup>+</sup> shape spaces were not significantly different for the first PC axis only. All other comparisons were not significantly different for PC1-PC4.

The ascending arm of the premaxilla is extremely short in 35 dpf AB fish, but elongates significantly by 100 dpf (Fig. 4). Premaxilla shape in 100 dpf TH- specimens was similar to that of newly ossified premaxillae in 35 dpf AB fish, in that they had very short ascending arms (Fig. 6). Both the maxillae and premaxillae in the upper jaws of TH- fish exhibited limited growth in size that was unaccompanied by the shape changes seen in those of post-metamorphic euthyroid and TH+ fish (Fig. 2 and 4).

Ascending arm length determines maximum upper jaw protrusion distance (Hulsey et al. 2010; Cooper, Carter, et al. 2017). Variation in ascending arm length was strongly associated with both post-metamorphic AB zebrafish development and danionin evolution (Fig. 4). Developmental variation in AB premaxilla shape was not significantly different from the variation in premaxilla shape that has evolved among nine additional danionin species (Table 3). Among the species that we examined, those from the genus *Danio* have longer premaxillary ascending arms than those from other danionin genera, with *Danio erythromicron* having the longest arms (Fig. 6). The premaxillae of *Danionella*, *Devario* and *Microdevario* were most similar to those from 35 dpf AB and 100 dpf TH- zebrafish. Manipulation of freshly euthanized specimens from these three genera, as well as observations of their feeding using high-speed video (pers. obs.; McMenamin, Carter, and Cooper 2017), indicate that jaw protrusion is extremely limited to non-existent in these species. This supports our fourth prediction that the functional morphology of jaw protrusion in adult TH- zebrafish will duplicate that of closely related minnows with limited protrusion abilities.

## DISCUSSION

We found strong evidence that TH levels affect the development of zebrafish feeding biomechanics (prediction 1), that the elimination of TH production reduces developmental variation in zebrafish head shape (prediction 2), and that the functional morphology of jaw protrusion in adult TH- zebrafish duplicates that of closely related minnows with limited protrusion abilities (prediction 4). Normal TH levels do not appear to be required for the development of the wild-type pattern of covariation between different regions of the zebrafish skull (prediction 3), but specimens with highly divergent TH levels (TH- and TH+) did exhibit limited differences in cranial covariation patterns (Table 3). We also found evidence that the developmental trajectory of premaxillary shape change in AB zebrafish is parallel to an important evolutionary axis of danionin diversification (premaxilla shape and jaw protrusion ability).

We found that differences in feeding biomechanics between our TH treatment groups were present at all of the developmental stages we examined, but that these differences increased with age and were particularly pronounced after metamorphosis (Table 1). The presence of TH appears to be particularly important for the transition from larval to adult feeding mechanics in the zebrafish, since a lack of TH delayed cranial ossification and arrested premaxillae formation before there was sufficient ascending arm development to permit upper jaw protrusion (Table 1; Fig. 2, 4 and 6; McMenamin, Carter, and Cooper 2017).

A loss of TH signaling appeared to truncate aspects of zebrafish cranial development. In addition to arresting premaxilla morphogenesis immediately after ossification (Fig. 4), lack of TH caused a significant reduction in the skull shape variation that arose between 8 and 100 dpf (Table 2). Conversely, TH+ fish showed an increase in developmental variation in skull shape relative to both DMSO and TH-, though they were not significantly different from AB specimens in this regard. Much of this increase in skull shape variation appeared to be due to the fact that high TH levels induced pronounced but variable mandibular prognathism in post-metamorphic fish (Fig. 2).

The only significant differences in cranial shape covariation that we observed were between the TH- and TH+ treatments, but these differences did not include PC1, which is the largest axis of shape variation (Table 3). Changes to patterns of anatomical trait covariation can have debilitating effects, especially in biomechanical systems where there is a high level of functional integration between different elements (Walker 2007; Armbruster et al. 2014; Kimmel et al. 2015). Both TH- and TH+ fish retained a sufficient level of integration between the various bones of the skull in order to feed successfully. The loss of TH shifted zebrafish development so that their adult premaxillary shape converged on those of other danionin species that feed without jaw protrusion (Fig. 4). Differences in jaw protrusion ability are strongly associated with differences in diet in marine damselfishes (Cooper, McGraw, et al. 2017), and although diet data for wild danionins is limited or non-existent, we suspect that the same may be true for the species examined here. We can therefore speculate that evolutionary changes in TH signaling may promote certain adaptive changes in fish skull mechanics without causing such a severe reorganization of cranial development as to produce non-functional skulls. It has been suggested that changes to TH signaling have played such a role during the adaptive diversification of cypriniform fishes (Shkil et al. 2012; Shkil et al. 2015; Shkil and Smirnov 2015; McMenamin, Carter, and Cooper 2017).

### The importance of late developmental periods to fish evo-devo

The field of evo-devo is focused on understanding the connections between developmental processes and evolutionary change (Carroll 2008). Phylogenetic analyses of comparative data have expanded tremendously in recent years and the evolutionary patterns traced by many lineages have been described in great detail (Freckleton et al. 2002; Garland et al. 2005; Mouquet et al. 2012). A particularly rich source of comparative information exists for the field of fish feeding biomechanics and we now know a great deal about the ecological, morphological and functional evolution of many fish clades. Due to the use of multiple fish species as model organisms for developmental study (e.g., zebrafish, medaka, Mexican tetra, fugu, stickleback, multiple African rift-lake cichlids, etc.) it is possible to experimentally explore aspects of fish development that have played important roles in shaping evolutionary diversification. However, since most evolutionary studies of fish feeding have focused on adult specimens and since most fishes do not develop adult feeding biomechanics until late in their development, often after a pronounced metamorphosis, merging these two areas of investigation will require an increased focus on the later stages of skull morphogenesis.

Evo-devo studies of jaw protrusion provide an illustration of this point. Protrusile jaws constitute one of the most significant biomechanical innovations to arise in fish skulls, but we know of no species in which protrusion arises before the larva to juvenile transition (Hernández, Barresi, and Devoto 2002; Power et al. 2008; McMenamin and Parichy 2013). Although there has been extensive study of skull morphogenesis in zebrafish embryos and larvae, investigation of these developmental periods has limited relevance to the development of protrusion ability. Thyroid hormone signaling has pervasive effects on skeletal morphogenesis during late development and our data support the assertion that a better understanding of how TH affects cranial remodeling has important relevance to the evo-devo of adaptive diversification in fish feeding.

## CONCLUSIONS

At the transition from larva to juvenile zebrafish, TH signaling induces development of an adult premaxilla shape and adult feeding kinematics. Hypothyroidism inhibited the development of jaw protrusion by severely reducing the development of premaxillary ascending arm length. Normal ontogenetic changes in zebrafish premaxillary shape represent developmental variation that could underlie evolutionary changes in jaw protrusion ability. The pronounced effects of the hormone on the development of the functional morphology of the skull during the juvenile to adult transition suggest that changes in TH or pathways affected by TH may have contributed to adaptive diversification of fish feeding biomechanics.

---

**Supplementary Figure 1.**
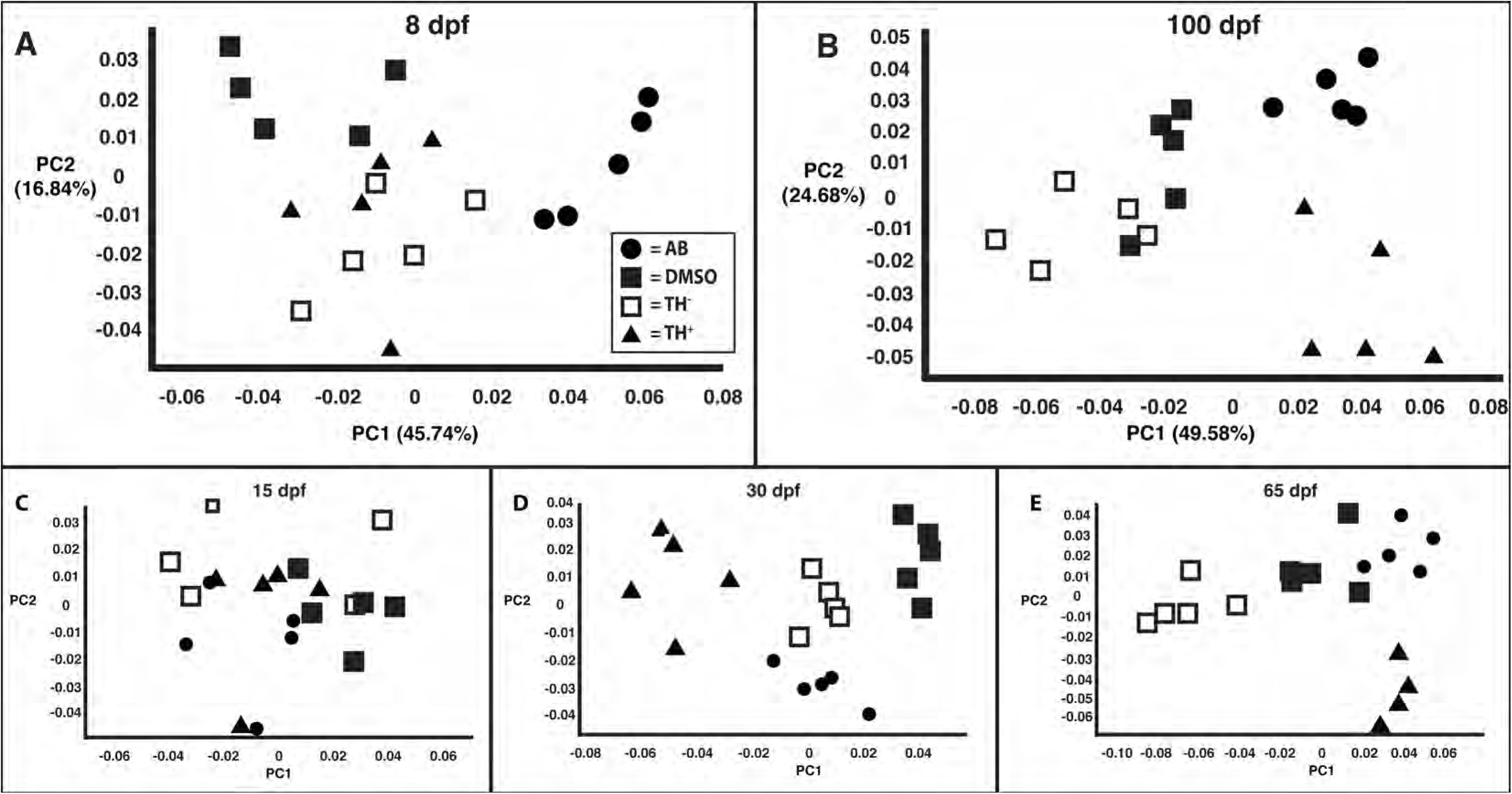
Comparison of the head shapes of the specimens used in kinematic analyses. Proximity denotes similarity. Key to symbols in panel A. A. Principal component score plot derived from head shape analyses of 8 dpf specimens. B. Principal component score plot derived from head shape analyses of 100 dpf specimens. C. Deformation grid and vector plot that shows the shape variation associated with PC1. D. Deformation grid and vector plot that shows the shape variation associated with PC2. E. Principal component score plot derived from head shape analyses of 15 dpf specimens. F. Principal component score plot derived from head shape analyses of 30 dpf specimens. G. Principal component score plot derived from head shape analyses of 65 dpf specimens.

**Supplementary Figure 2.**
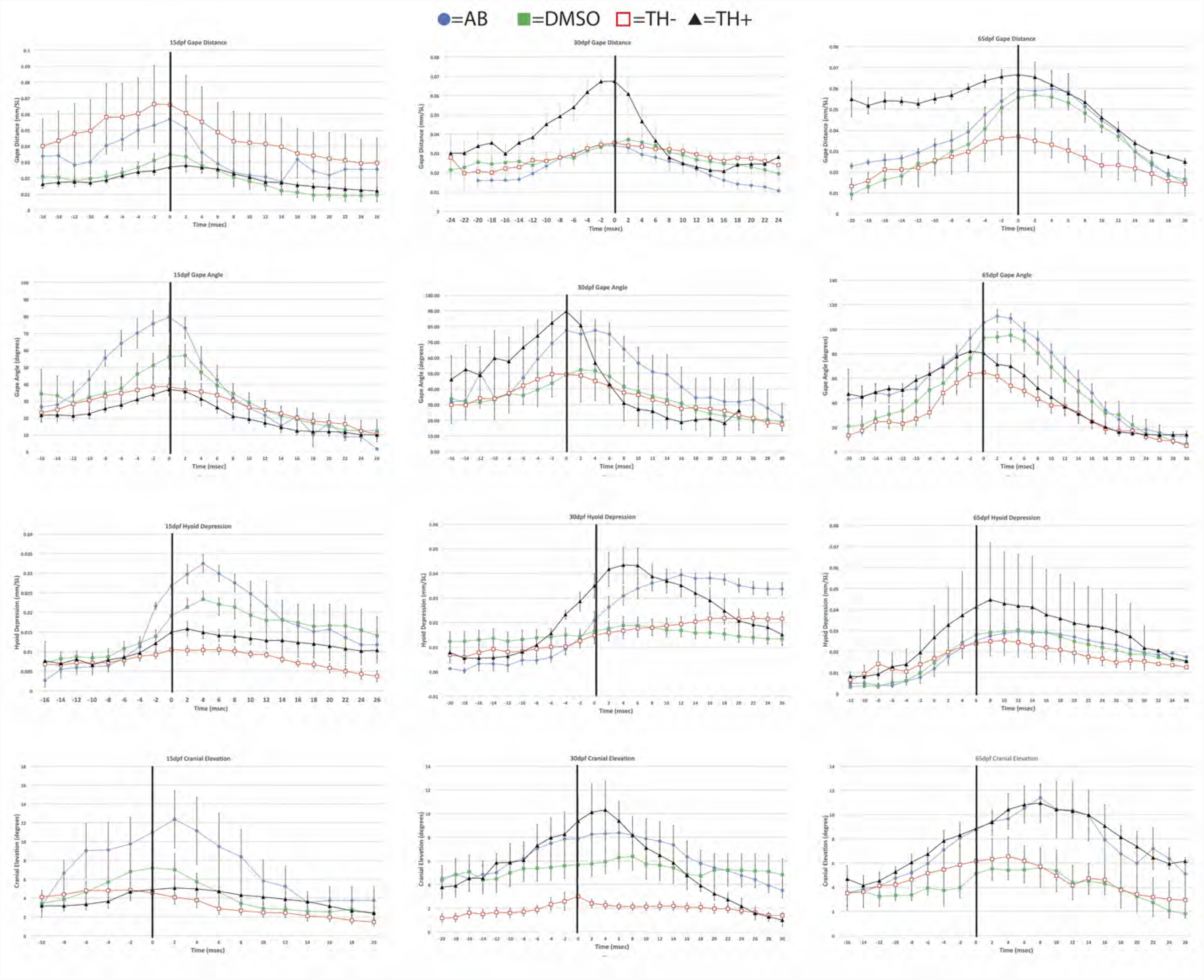
Comparisons of cranial movements during feeding among the four TH treatment groups at 15, 30 and 65 dpf. In all cases time zero represents the time point when live prey (paramecia at 15 dpf and brine shrimp nauplii at 30 and 65 dpf) passed the tips of the upper and lower jaws as they were being engulfed. Units and standardizations are given in parentheses in each case. SL = standard length.

## REFERENCES

Albertson, R. C. and Yelick, P. C. 2004. Morphogenesis of the jaw: Development beyond the embryo. In Zebrafish: 2nd Edition Cellular and Developmental Biology. San Diego: Elsevier Academic Press Inc.

Armbruster, W. S., Pelabon, C., Bolstad, G. H., and Hansen, T. F. 2014. Integrated phenotypes: understanding trait covariation in plants and animals. Philosophical Transactions of the Royal Society B-Biological Sciences 369 (1649):16.

Carroll, S. B. 2008. Evo-devo and an expanding evolutionary synthesis: a genetic theory of morphological evolution. Cell 134.

Cooper, K., McGraw, A., Khazanchi, D., and Ieee. 2017. Bioinformatics for Middle School Aged Children: Activities for Exposure to an Interdisciplinary Field. In Proceedings of the 2017 7th Ieee Integrated Stem Education Conference.

Cooper, W. J., Carter, C. B., Conith, A. J., Rice, A. N., and Westneat, M. W. 2017. The evolution of jaw protrusion mechanics is tightly coupled to bentho-pelagic divergence in damselfishes (Pomacentridae). The Journal of Experimental Biology 220 (4):652–666.

Cooper, W. J., Parsons, K., McIntyre, A., Kern, B., McGee-Moore, A., and Albertson, R. C. 2010. Bentho-Pelagic Divergence of Cichlid Feeding Architecture Was Prodigious and Consistent during Multiple Adaptive Radiations within African Rift-Lakes. PLoS ONE 5 (3):A38–A50.

Cooper, W. J., Wernle, J., Mann, K. A., and Albertson, R. C. 2011. Functional and Genetic Integration in the Skulls of Lake Malawi Cichlids. Evolutionary Biology 38 (3):316–334.

Cooper, W. J. and Westneat, M. W. 2009. Form and function of damselfish skulls: rapid and repeated evolution into a limited number of trophic niches. BMC Evolutionary Biology 9.

Cooper, W. J., Wirgau, R. M., Sweet, E. M., and Albertson, R. C. 2013. Deficiency of zebrafish fgf20a results in aberrant skull remodeling that mimics both human cranial disease and evolutionarily important fish skull morphologies. Evolution & Development 15 (6):426– 441.

Das, B., Cai, L. Q., Carter, M. G., Piao, Y. L., Sharov, A. A., Ko, M. S. H., and Brown, D. D. 2006. Gene expression changes at metamorphosis induced by thyroid hormone in Xenopus laevis tadpoles. Developmental Biology 291 (2):342–355.

Desjardin, C., Charles, C., Benoist-Lasselin, C., Riviere, J., Gilles, M., Chassande, O., Morgenthaler, C., Laloe, D., Lecardonnel, J., Flamant, F., Legeai-Mallet, L., and Schibler, L. 2014. Chondrocytes Play a Major Role in the Stimulation of Bone Growth by Thyroid Hormone. Endocrinology 155 (8):3123–3135.

Ferry-Graham, L. A., Gibb, A. C., and Hernandez, L. P. 2008. Premaxillary movements in cyprinodontiform fishes: An unusual protrusion mechanism facilitates “picking” prey capture. Zoology 111 (6):455–466.

Foote, M. 1993. Contributions of Individual Taxa to Overall Morphological Disparity. Paleobiology 19 (4):403–419.

Freckleton, R. P., Harvey, P. H., and Pagel, M. 2002. Phylogenetic analysis and comparative data: A test and review of evidence. American Naturalist 160 (6):712–726.

Garland, T., Bennett, A. F., and Rezende, E. L. 2005. Phylogenetic approaches in comparative physiology. Journal of Experimental Biology 208 (16):3015–3035.

Hanken, J. and Hall, B. K. 1988. Skull development during anuran metamorphosis. 2. role of thyroid-hormone in osteogenesis. Anatomy and Embryology 178 (3):219–227.

Hanken, J. and Summers, C. H. 1988. Skull development during anuran metamorphosis. 3. role of thyroid-hormone in chondrogenesis. Journal of Experimental Zoology 246 (2):156– 170.

Hernandez, L. P. 1995. The functional morphology of feeding in three ontogenetic stages of the zebrafish, Danio rerio. American Zoologist 35 (5):104A–104A.

Hernandez, L. P. 2000. Intraspecific scaling of feeding mechanics in an ontogenetic series of zebrafish, Danio rerio. Journal of Experimental Biology 203 (19):3033–3043.

Hernández, L. P., Barresi, M. J. F., and Devoto, S. H. 2002. Functional morphology and developmental biology of zebrafish: Reciprocal illumination from an unlikely couple. Integrative and Comparative Biology 42 (2):222–231.

Hirano, A., Akita, S., and Fujii, T. 1995. Craniofacial deformities associated with juvenile hyperthyroidism. Cleft Palate-Craniofacial Journal 32 (4):328–333.

Holzman, R., Collar, D. C., Mehta, R. S., and Wainwright, P. C. 2012. An integrative modeling approach to elucidate suction-feeding performance. Journal of Experimental Biology 215 (1):1–13.

Hulsey, C. D., Hollingsworth, P. R., Jr., and Holzman, R. 2010. Co-evolution of the premaxilla and jaw protrusion in cichlid fishes (Heroine: Cichlidae). Biological Journal of the Linnean Society 100 (3):619–629.

Kimmel, C. B., Watson, S., Couture, R. B., McKibben, N. S., Nichols, J. T., Richardson, S. E., and Noakes, D. L. G. 2015. Patterns of variation and covariation in the shapes of mandibular bones of juvenile salmonids in the genus Oncorhynchus. Evolution & Development 17 (5):302–314.

Konow, N. and Bellwood, D. R. 2005. Prey-capture in Pomacanthus semicirculatus (Teleostei, Pomacanthidae): functional implications of intramandibular joints in marine angelfishes. Journal of Experimental Biology 208 (8):1421–1433.

Laudet, V. 2011. The Origins and Evolution of Vertebrate Metamorphosis. Current Biology 21 (18):R726–R737.

Leis, J. M. and McCormick, M. I. 2006. The biology, behavior and ecology of the pelagic, larval stage of coral reef fishes. In Coral reef fishes: Dynamics and diversity in a complex ecosystem, edited by P. F. Sale. Burlington, MA: Elsevier, Inc.

Martinez-Abadias, N., Esparza, M., Sjovold, T., Gonzalez-Jose, R., Santos, M., and Hernandez, M. 2009. Heritability of human cranial dimensions: comparing the evolvability of different cranial regions. Journal of Anatomy 214 (1):19–35.

McCluskey, B. M. and Postlethwait, J. H. 2015. Phylogeny of Zebrafish, a “Model Species,” within Danio, a “Model Genus”. Molecular Biology and Evolution 32 (3):635–652.

McCormick, M. I., Makey, L., and Dufour, V. 2002. Comparative study of metamorphosis in tropical reef fishes. Marine Biology 141 (5):841–853.

McCormick, M. I. and Makey, L. J. 1997. Post-settlement transition in coral reef fishes: overlooked complexity in niche shifts. Marine Ecology Progress Series 153:247–257.

McMenamin, S., Carter, C., and Cooper, W. J. 2017. Thyroid Hormone Stimulates the Onset of Adult Feeding Kinematics in Zebrafish. Zebrafish 14 (6):517–525.

McMenamin, S. K., Bain, E. J., McCann, A. E., Patterson, L. B., Eom, D. S., Waller, Z. P., Hamill, J. C., Kuhlman, J. A., Eisen, J. S., and Parichy, D. M. 2014. Thyroid hormone–dependent adult pigment cell lineage and pattern in zebrafish. Science 345 (6202):1358–1361.

McMenamin, S. K. and Parichy, D. M. 2013. Metamorphosis in Teleosts. In Animal Metamorphosis, edited by Y. B. Shi. San Diego: Elsevier Academic Press Inc.

Mouquet, N., Devictor, V., Meynard, C. N., Munoz, F., Bersier, L. F., Chave, J., Couteron, P., Dalecky, A., Fontaine, C., Gravel, D., Hardy, O. J., Jabot, F., Lavergne, S., Leibold, M., Mouillot, D., Munkemuller, T., Pavoine, S., Prinzing, A., Rodrigues, A. S. L., Rohr, R. P., Thebault, E., and Thuiller, W. 2012. Ecophylogenetics: advances and perspectives. Biological Reviews 87 (4):769–785.

Near, T. J., Dornburg, A., Eytan, R. I., Keck, B. P., Smith, W. L., Kuhn, K. L., Moore, J. A., Price, S. A., Burbrink, F. T., Friedman, M., and Wainwright, P. C. 2013. Phylogeny and tempo of diversification in the superradiation of spiny-rayed fishes. Proceedings of the National Academy of Sciences.

Paris, M., Escriva, H., Schubert, M., Brunet, F., Brtko, J., Ciesielski, F., Jamin, E., Cravedi, J. P., Renaud, J. P., Scanlan, T. S., Holland, N. D., and Laudet, V. 2010. Amphioxus thyroid hormone signaling pathway and the evolution of metamorphosis in chordates. Integrative and Comparative Biology 50:E133–E133.

Parsons, K., Andreeva, V., Cooper, W. J., Yelick, P. C., and Albertson, R. C. 2010. Morphogenesis of the zebrafish jaw: Development beyond the embryo. In *Methods in Cell Biology: The Zebrafish*, edited by M. Westerfield, H. W. Detrich and L. I. Zon. San Diego: Elsevier Academic Press Inc.

Pothoff, T. 1984. Clearing and staining techniques. In Ontogeny and Systematics of Fishes, edited by H. G. Moser, W. J. Richards, D. M. Cohen, M. P. Fahay, A. W. Kendall Jr. and S. L. Richardson. Lawrence, Kansas: Allen Press.

Power, D. M., Silva, N., and Campinho, M. A. 2008. Metamorphosis. In Fish Larval Physiology, edited by R. N. Finn and B. G. Kapoor. Enfield, New Hampshire: Science Publishers.

Shkil, F. N., Kapitanova, D. V., Borisov, V. B., Abdissa, B., and Smirnov, S. V. 2012. Thyroid hormone in skeletal development of cyprinids: effects and morphological consequences. Journal of Applied Ichthyology 28 (3):398–405.

Shkil, F. N., Lazebnyi, O. E., Kapitanova, D. V., Abdissa, B., Borisov, V. B., and Smirnov, S. V. 2015. Ontogenetic mechanisms of explosive morphological divergence in the Lake Tana (Ethiopia) species flock of large African barbs (Labeobarbus; Cyprinidae; Teleostei). Russian Journal of Developmental Biology 46 (5):294–306.

Shkil, F. N. and Smirnov, S. V. 2015. Experimental approach to the hypotheses of heterochronic evolution in lower vertebrates. Paleontological Journal 49 (14):1624–1634.

Simon, M. N., Machado, F. A., and Marroig, G. 2016. High evolutionary constraints limited adaptive responses to past climate changes in toad skulls. Proceedings of the Royal Society B-Biological Sciences 283 (1841).

Staab, K. L., Holzman, R., Hernandez, L. P., and Wainwright, P. C. 2012. Independently evolved upper jaw protrusion mechanisms show convergent hydrodynamic function in teleost fishes. Journal of Experimental Biology 215 (9):1456–1463.

Tang, K. L., Agnew, M. K., Hirt, M. V., Sado, T., Schneider, L. M., Freyhof, J., Sulaiman, Z., Swartz, E., Vidthayanon, C., Miya, M., Saitoh, K., Simons, A. M., Wood, R. M., and Mayden, R. L. 2010. Systematics of the subfamily Danioninae (Teleostei: Cypriniformes: Cyprinidae). Molecular Phylogenetics and Evolution 57 (1):189–214.

Wainwright, P. C., McGee, M. D., Longo, S. J., and Hernandez, L. P. 2015. Origins, Innovations, and Diversification of Suction Feeding in Vertebrates. Integrative and Comparative Biology 55 (1):134–145.

Walker, J. A. 2007. A general model of functional constraints on phenotypic evolution. American Naturalist 170 (5):681–689.

Wojcicka, A., Bassett, J. H. D., and Williams, G. R. 2013. Mechanisms of action of thyroid hormones in the skeleton. Biochimica Et Biophysica Acta-General Subjects 1830 (7):3979–3986.

Yang, L., Mayden, R. L., Sado, T., He, S. P., Saitoh, K., and Miya, M. 2010. Molecular phylogeny of the fishes traditionally referred to Cyprinini sensu stricto (Teleostei: Cypriniformes). Zoologica Scripta 39 (6):527–550.

Yaniv, S., Elad, D., and Holzman, R. 2014. Suction feeding across fish life stages: flow dynamics from larvae to adults and implications for prey capture. Journal of Experimental Biology 217 (20):3748–3757.

